# Unravelling a diversity of cellular structures and aggregation dynamics during the early development of *Myxococcus xanthus*

**DOI:** 10.1101/2024.07.01.601548

**Authors:** Natsuko Rivera-Yoshida, Alejandro V. Arzola, Mariana Benítez

## Abstract

Aggregation underlies collective dynamics of a diversity of organisms, enabling the formation of complex structures and emergent behaviours in interaction with the environment. *Myxococcus xanthus*, a social bacterium, is a valuable model for studying aggregation, particularly in the development and evolution of multicellularity. While the collective behaviour of *M. xanthus* has been largely studied under very crowded, ideal, conditions, there is a lack of understanding at not crowded or sparse conditions. In this work, we study the early stages of emergent collective behaviour of *M. xanthus* under nutrient-poor and low density conditions, uncovering the formation of diverse cellular structures with different shapes and sizes, ranging from individual cells to networks comprising thousands of cells. We study their motility patterns and their prevalence along development, and argue over their cross-scale role on the population’s exploratory dynamics. This work contributes to understanding key, yet largely understudied, aspects in the early stages of multicellular development in myxobacteria, shedding light on the dynamics underlying aggregative processes in this and other taxa and study systems.

## INTRODUCTION

Aggregation is a ubiquitous phenomenon of non-equilibrium systems that leads to the emergence of spatial structures both in living and non-living matter (Ginot et al., 2018; Arias Del Angel et al., 2020; Fu et al., 2022; Ben et al., 2023; Newman, 2024). In particular, aggregation among self-propelled particles in the microscopic world arises from their diverse attractive interactions and confinement, exhibiting structures that vary in size and shape across spatio-temporal scales (Vernerey et al., 2019). Cellular aggregation constitutes one of the routes to collective motility and multicellular development, and understanding it can shed light on the ecological and evolutionary processes underlying the formation of biofilms and multicellular structures in diverse prokaryote and eukaryote lineages (Niklas and Newman, 2016; Herron et al., 2022). *Myxococcus xanthus*, a motile rod-shaped bacterium that glides across surfaces, exhibits different collective behaviours under nutrient-availability conditions. Under starvation, *M. xanthus* cells undergo aggregation through a developmental process that can culminate in the formation of differentiated multicellular structures known as fruiting bodies (Whitworth, 2008; Zhang et al., 2012; Muñoz-Dorado et al., 2016). Because of this, *M. xanthus* serves as a valuable model system to study the aggregative way to multicellularity, a major transition in the evolutionary history of life (Arias Del Angel et al., 2017, 2020; Zhang et al, 2011).

*M. xanthus* aggregation exhibits a remarkable richness of cellular structures and behaviours depending in part on the cellular density (Peruani et al., 2012; Thutupalli, et al., 2015; Liu et al., 2019). These have been well-studied at high, standard laboratory densities, at which cells form dense mats that completely cover the substrate, tend to remain in contact and align in a regime that mimics a nematic liquid (Thutupalli et al., 2015). Aggregation dynamics are also driven by cellular reversal timing and correlated with the nearest aggregate size, which, in turn, correlates with local cell density (Thutupalli et al., 2015, Murphy et al., 2023). In a *M. xanthus* mutant that is unable to reverse and has been studied at nutrient-rich conditions, collective motion emerges only above a density value allowing the formation of large structures (Peurani, 2012). Interestingly, isolated cells remain non-motile but can be induced into motion by motile cells (Hodgkin and Kaiser, 1977). However, there is no systematic account of how *M. xanthus* cells at low densities and in the developmental regime (nutrient-poor conditions) organise and transit from single cells to cellular structures, especially during the early stages of development when cells are beginning to interact with themselves and the substrate. Addressing this gap may uncover aspects of aggregation that are impossible to detect at higher cellular densities but might be critical at the onset of development.

In this study, we describe and analyse the organisation of sparse populations of *M. xanthus* cells in nutrient-poor and low packing density conditions during the initial 8 h of development. Our experimental setup allowed us to acquire image and video for large areas of up to 1000 µm x 624 µm. We explore how cellular organisation changes with cellular density and developmental time. With this, we contribute to unpacking and understanding largely understudied stages and processes in the development of multicellular structures in myxobacteria, and to further understanding aggregation as an ubiquitous phenomenon in nature.

## METHODS

### Experimental growth and developmental conditions

Following Yang and Higgs (2014), DK1622 strain was taken from a frozen stock by spotting 50 µl onto a casitone yeast extract (CYE) agar plate [1% Bacto Casitone, 10 mM tris-HCl (pH 7.6), 0.5% yeast extract, 10 mM Mops (pH 7.6), and 4 mM MgSO4; BD Bacto™ Agar] and incubated at 32°C for 2 days. Cells from the resulting colony were transferred to 25 ml of CYE liquid medium and incubated at 32°C, shaking at 250 rpm overnight. The culture was diluted to 0.7 OD550 (nutrient-rich liquid culture).

For the development assays, cells were harvested by spinning them at 8000 rpm for 5 min. The pellet was washed with a tris phosphate magnesium (TPM) medium solution [10 mM tris-HCl (pH 7.6), 1 mM K2HPO4, and 8 mM MgSO4] and resuspended in 1/10 of the original volume. This cellular stock was diluted 1:350, 1:35 and 1:10 corresponding to approximately 0.002, 0.02 and 0.07 OD550. Fifteen microliters of each dilution were spotted onto 5 ml TPM agar plates (BD Bacto™ Agar, 2.4%) and incubated at 28°C for 8 h. The essays were repeated three times for statistical support. When spotted and dried onto agar plates, the cells in the three dilutions exhibit 0.02, 0.1, 0.23 packing densities, respectively, showing a qualitatively different organisation: mostly isolated single cells, crowded single-cells and some pairs of cells, and sparse small groups of cells, respectively. We will refer to these conditions as low, medium and high density, although they are all relatively low in comparison with the densities usually used in *M. xanthus* essays.

### Microscopic local observation, image and video acquisition and analysis

The samples were observed through the agar growing medium using an inverted optical microscope with a 10X objective lens (Olympus UPlanSApo series, 0.4 NA), illuminated with amber light and recorded with a Basler acA 1600-20uc with overall resolution of 0.52 µm/pixel and 1000 µm x 624 µm field of vision. The microscope was set inside an isolated chamber with a controlled temperature of 28 C. Micrographs were taken for two of the three replicates, while 15 min videos were taken at t=0, 4 and 8 h from the onset of starvation (after the spots dried) for the third replica at an acquisition rate of 1 frame per second. Micrographs and video frames were binarized using segmentation through edge-detection and intensity-threshold criteria implemented with the Image Processing Toolbox of MATLAB. Analysis from the resulting database was conducted using R v.4.2.0.

To study the exploration of space at the population level, we defined the fraction of explored area as 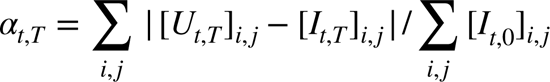, with 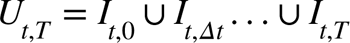 the union of the binarized images, from the developmental time t to t+T taken every Δt=50 s. Fig. 3E shows *α_t,T_* as a function of the three developmental times (t=0, 4 and 8 h) and T=600 s for cultures with medium and high cell density (cells at low density and t=0 s practically do not explore new areas). The resulting plots reflect the overall changes due to the change in the position, shape and size of the different cellular structures over time.

## RESULTS

### Diverse cellular structures dynamically arise and can coexist along development

To explore the effect of cellular density on the early developmental dynamics of *M. xanthus*, we followed the dynamics of cellular structures in the wild-type (WT)-DK1622 strain at three different cellular densities (all of them relatively low, but referred here as low, medium and high (Methods)). Micrographs and 15-minute videos were taken at 0, 4, and 8 h for each condition. Through this time lapse, we monitored the formation and eventual coexistence of motile structures exhibiting different morphologies and a wide range of sizes, and dynamically forming through continuous merging and dissociation of individual cells and other cellular structures (Figure 1A). We first describe qualitatively these structures in terms of their size in number of cells (ranging from one to thousands, and from a scale of tens to hundreds of microns), their spatial exploration mode (directed or not), presence of reversals and the stability of their boundaries (see a more detailed and quantitative description in Fig. 3).

**Figure 1.**
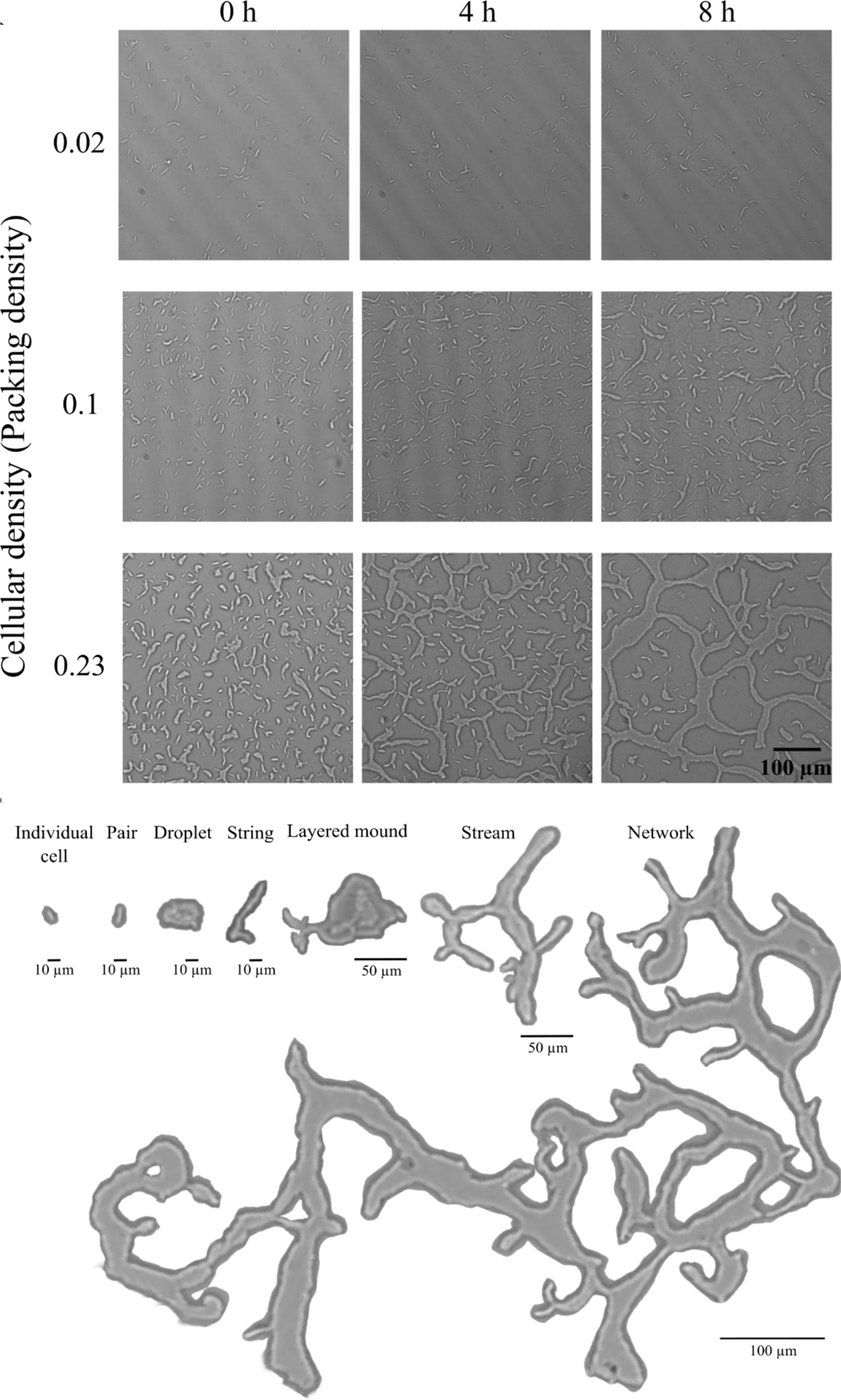
*M. xanthus* early development (from 0 to 8 h) with varying cellular densities (packing densities). (A) At low cellular density, *M. xanthus* cells stay as single-cells, pairs or triads through time. At medium cellular density, single-cells and small elongated cellular structures coexist in dynamic arrangements in a homogeneous spatial distribution. At high cellular density, single-cells and larger cellular structures varying their size and shape, coexist in dynamic arrangements. (B) Typology of structures with different sizes. Morphologies and behaviours emerge through the continuous merging and dissociating of cells and groups of cells, coexisting along development.

Considering these features, we propose a typology of cellular structures that applies for *M. xanthus* early development at low densities, but is not restricted to these conditions (Figure 1B, Supplementary videos 1-9). *Individual cells* tend to move locally, without an apparent target or large net displacement, oscillating back and forth along the trails they form. When arranged in *pairs* through head-to-coil contacts their motility is qualitatively similar to that of individual cells, but they can come together and split as the cells in the pair reverse. *Strings* are elongated, thin and flexible structures characterised by unstable boundaries and conformed by 3 to tens of cells. These structures can maintain their elongated form or transit into more rounded structures. *Droplets* include a diversity of irregular but overall rounded morphological structures that can include up to hundreds of cells and exhibit rapid and persistent movement with an apparent target, often merging and dissolving rapidly upon contact with individual cells and other cellular structures. *Layer mounds* are rounded structures composed of two or more layers that can move as a single structure, though the layers can also move independently. Droplets may coalesce into layered mounds or wide branched *streams*, each comprising hundreds of cells and exhibiting also unsteady boundaries, but different in their morphology and behaviour from droplets and strings (Figure 1B). Within streams, cells align and tend to move collectively, with anisotropic motion. *Networks* are branched and large structures that can encompass almost all cells within a sample. While these networks exhibit continuous shape changes and single-cells within move (typically aligned), they maintain relatively stable boundaries over extended periods. Overall, cellular structures seem to move more rapidly than individual cells, exploring larger areas of the substrate. While some of these structures and phenomena have been previously described from an active matter perspective, e.g. cell flocks and streams in Thutupalli and coworkers (2015), or rafts and peninsulas in the study of cellular motility in the edge of a swarm by Kaiser and Crosby (1983), our proposed typology highlights a variety of previously unreported cellular structures and behaviours involved in the early aggregation dynamics of *M. xanthus*.

### The prevalence of cellular structures changes along development and with varying initial densities

We observe that the prevalence of different cellular structures and the overall cellular organisation change over time and with initial cellular density (Figure 2). To better understand how these changes occur, we quantified the size and number of the cellular structures present at 0, 4 and 8 h.

**Figure 2.**
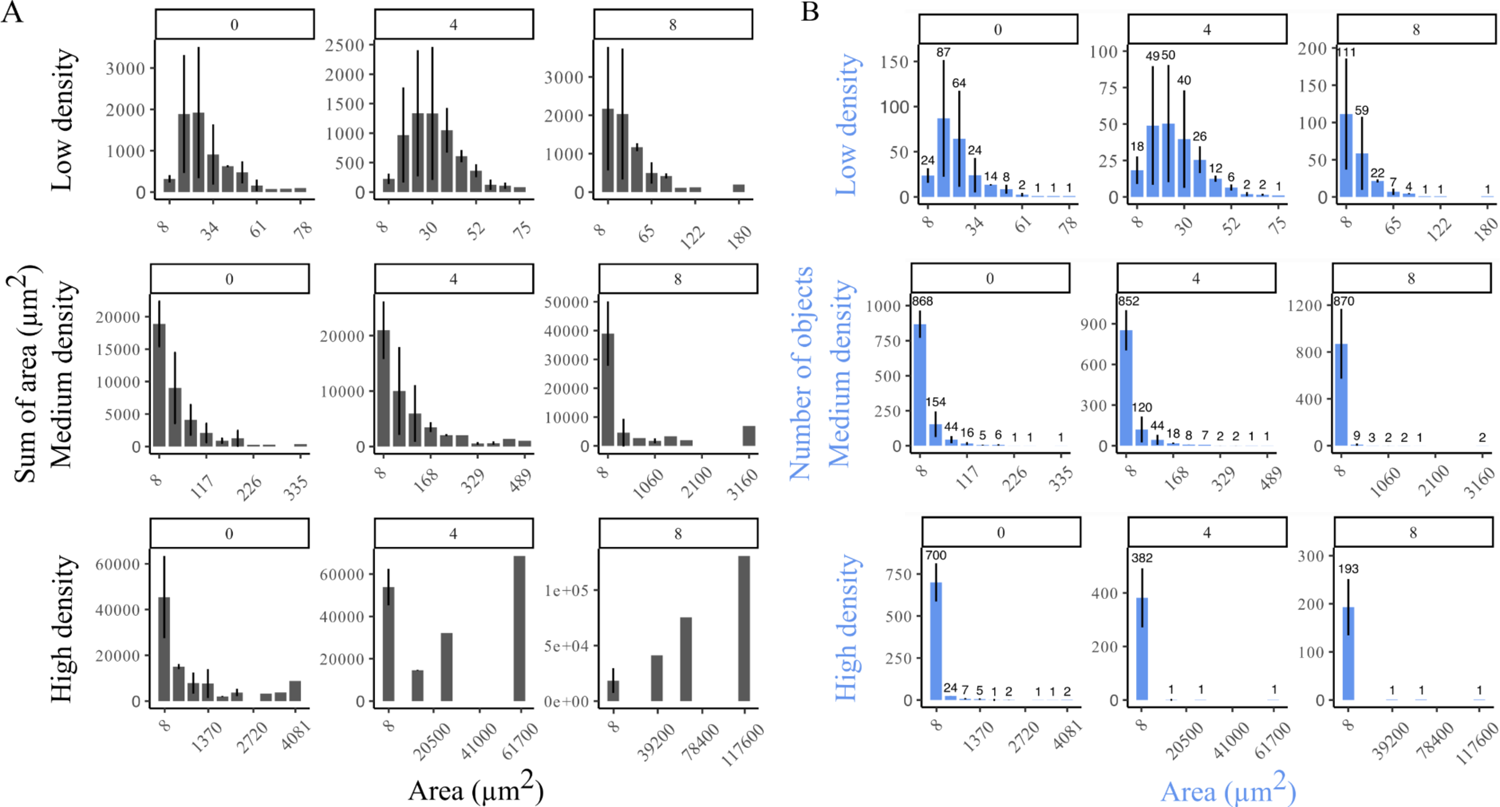
Occupied area and frequency of structure sizes for different initial density conditions throughout early development. (A) Area occupied and (B) number of objects for different structure sizes. Data was obtained from 3 replicates.

At low cellular density, individual cells and pairs were the dominant structures (Fig. 2A, 2B). At medium cellular density, small structures prevailed as individual cells, pairs and strings, coexisting at 0 h. Even if structures were in general small in the medium density condition, a couple of strings occupying a significant area form at 8 h (Fig 1A, 2A, 2B). At high cellular density, the presence of individual cells decreased through time. Small groups of cells were the most frequent structures, although small structures for this condition were much larger than small structures in low and medium density conditions, corresponding mostly to strings, droplets and streams (Figure 2A, 2B). In the high density condition, the morphology of large structures also varies drastically through time. It starts from droplets, streams and layered mounds at 0 h, then these merge and dissolve into and from branched streams and droplets at 4 h, and at 8 h end up forming a single network that occupies an area five orders of magnitude larger that the small structures in the sample (Figure 1; Figure 2A, 2B; Videos 7-9).

### Different types of cellular structures promote different ways of motility at the structure and population levels

In order to explore how the prevalence of different structures impacts cellular dynamics across levels of organisation, we further describe the structures’ dynamics and quantify the area explored by cells and structures at the whole population level. Small to medium-sized structures can move as a whole over empty space (e.g., the groups indicated by yellow arrows in Figs. 3A-B) or elongate in different directions, stretching and breaking into parts (green arrows). In contrast, large networks tend to confine the dynamics of cells, resulting in more stable structures, even though their shape is constantly changing due to spontaneously growing zones (indicated by yellow arrows in Figs. 3C-D) or stretching zones (indicated by green arrows). As expected from the description above, high cell density populations exhibit lower values of the fraction of explored area (*α_t,T_*) than those at medium cell density. For medium cell density populations, *α_t,T_* shows a similar value for 4 and 8 h, while for the high cell density populations, there is an abrupt decay at 8 h. This aligns with the morphological changes observed at these two times, suggesting that large networks constrain cellular exploration. Overall, the behaviour of the cellular structures modifies the way in which cells explore the space and thus, the population patterns.

**Figure 3.**
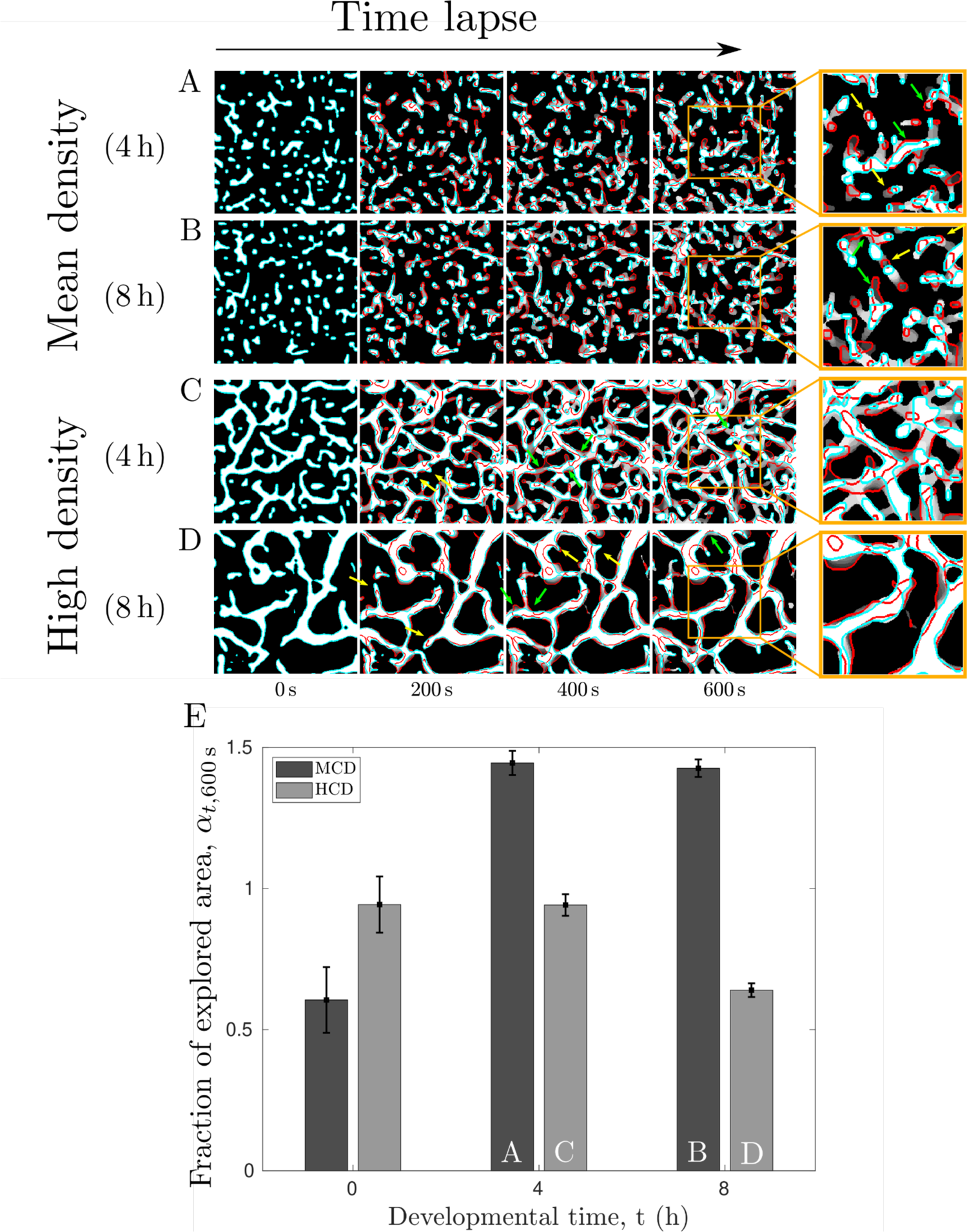
Dynamics of cellular structures at the population scale. (A-D) Medium and high cellular densities through the time lapse, at developmental times t= 4 and 8 h. The images at the first column show the cell structures in white delimited with cyan at the time lapse 0 s. These initial structures vary in shape, size and position through time, as illustrated in the second, third and fourth columns corresponding to time lapses 200, 400 and 600 s. The structures at each given time are shown in white delimited by cyan and the initial cellular arrangement at 0 h is delimited by red. Grey scale traces illustrate the gradual shape transition from 0 s to the time lapse, going from darker values at earlier stages to white at the final time point. Figures at the rightmost column correspond to an enlarged small section of the figure at 600 s. Yellow and green arrows in (A-B) indicate the motility pattern of small and medium size groups, while in (C-D) indicate growing and stretching zones. (E) Fraction of explored area (see the definition of *α_t,T_* in the main text) along T=600 s for the mean and high cell densities at t= 0, 4 and 8 h. Capital letters (A, C, B, D) in the columns indicate the corresponding conditions depicted in figures above.

## DISCUSSION AND CONCLUSIONS

We describe the cellular structures that arise dynamically during the early stages of aggregation in *M. xanthus* development at low cellular densities (Figure 1). These structures exhibit an ample range of sizes, morphologies, behaviours and boundary stability. By quantifying the prevalence of these structures at different times, we uncover sequences and transitions in the prevalence of specific structures (Figures 2 and 3). We also show how the sequences and transitions of different structures depend on cellular density. Notably, changes in the prevalence of cellular structures are associated with changes in patterns and motility at the population scale (Figure 3).

We propose a typology for the different cellular structures we observe (Figure 1). It is important to acknowledge that these structures are not static or discrete entities but rather dynamic, constantly changing in size and shape, often merging with or dissociating from other structures. Although ephemeral structures tend to be substituted by larger or more permanent ones, the transitions between them are density-dependent and do not strictly follow a determined sequence (Figures 2 and 3). Moreover, small and large structures can coexist for the whole time lapse (Figure 3B).

Studying *M. xanthus* behaviour and aggregation at low densities allowed us to characterise understudied types of structures such as “networks”, “strings” or “droplets”, which might be relevant to bacterial dispersion or aggregation in diverse ecological contexts. Additionally, our results challenge the notion that cells always tend to organise, as an intrinsic objective, into larger and steadier aggregates; they shift back and forth among structures of varying sizes and durability before aggregating into large steady structures, when they do. Our results align with previous evidence showing that, even at higher densities, not all cells form steady aggregates, but some disperse instead (Murphy et al., 2023; Rossine et al., 2020); that for extremely low densities, isolated cells rarely move or aggregate (Hodgkin and Kaiser, 1977); and that myxobacteria cells stimulate and affect each other’s movement via contact receptors, capillary action, stigmergy and fibrils (Dworkin et al., 1983; Gloag et al., 2016; Ramos et al., 2021; Behmlander and Dworkin, 1991). Altogether, our findings emphasise that starvation or stress are not a sufficient condition to trigger aggregation, but that cell density and probably other physical and cellular environmental conditions have to be met, or produced by cells themselves.

Moreover, our experimental design allowed us to test the effect of initial densities on structure formation and aggregation dynamics. Changes in cellular density led to qualitative changes in aggregation trajectories and the prevalence of the different types of cellular structures. This holds ecological significance because the density of *M. xanthus* in natural environments can be much lower that densities usually used in laboratory conditions, and can vary with substrate textures, topography, liquid content, nutrient availability, among other biological and physicochemical factors (Velicer et al., 1998; Kadam & Velicer, 2006; Rivera-Yoshida et al., 2020; H. Ramos et al., 2021). Cellular density within a population is often heterogeneous, as both physical and cellular environmental conditions fluctuate through time and space as cells move and organise into different structures. Indeed, local density appears to be central in determining the behaviour of cells and groups of cells (Murphy et al. 2023), but its effect on groups of different structures remains to be further unexplored.

Our findings regarding the role of cellular density can help us further understand the ecological conditions, including population density, in which aggregative multicellularity may have emerged in the evolutionary history of life. They can also help us understand how it may continue to arise when challenged by external stimuli or in interaction with different environments, including clinically relevant ones. Our results can indeed inform and dialogue with clinical studies aiming to manage infection control and treatment of bacterial populations by understanding the coexistence and development of spatial and temporal structures during biofilm formation (Bidossi et al., 2020; Knott et al., 2021; Rivera-Yoshida et al., 2022).

In conclusion, our experiments and analyses allowed us to uncover and begin to understand a wide range of cellular structures that arise during the aggregative development of *M. xanthus*, further unravelling specific aspects of this route to development and contributing to a nuanced understanding of the branched and diverse trajectories in which aggregation can occur in different cellular and ecological conditions, not only in *M. xanthus* but also in other aggregative multicellular groups.

## ACKNOWLEDGEMENTS

We thank Blanca Hernández Bautista, Blanca Hernández Hernández and Corina H. Ramos for technical and administrative assistance; and members of Laboratorio de Micromanipulación Óptica and LaParcela for insightful discussions. The authors gratefully acknowledge the financial support of the John Templeton Foundation (#62220). The opinions expressed in this paper are those of the authors and not those of the John Templeton Foundation.

## DATA ACCESSIBILITY

The database was uploaded as supplementary material in the submission system.

## SUPPLEMENTARY VIDEOS

Videos can be downloaded here (we suggest to play them at 4X or 8X): https://wetransfer.com/downloads/3d9ff61c99c539330e3e3d6af8b7a6f820240212051827/6af03d4e0e13eeac9db83acf0eab79da20240212051827/2b47a7

